# High Frequencies of Phenotypically and Functionally Senescent and Exhausted CD56^+^CD57^+^PD-1^+^ Natural Killer Cells, SARS-CoV-2-Specific Memory CD4^+^ and CD8^+^ T cells Associated with Severe Disease in Unvaccinated COVID-19 Patients

**DOI:** 10.1101/2022.07.26.501655

**Authors:** Ruchi Srivastava, Nisha Dhanushkodi, Swayam Prakash, Pierre Gregoire Coulon, Hawa Vahed, Latifa Zayou, Afshana Quadiri, Lbachir BenMohamed

**Affiliations:** Laboratory of Cellular and Molecular Immunology, Gavin Herbert Eye Institute, University of California Irvine, School of Medicine, Irvine, CA 92697; Department of Molecular Biology & Biochemistry; TechImmune, LLC, University Lab Partners, Irvine, CA 92660-7913; Department of Vaccines and Immunotherapies, TechImmune, LLC, University Lab Partners, Irvine, CA 92660-7913; Institute for Immunology; University of California Irvine, School of Medicine, Irvine, CA 92697

**Keywords:** SARS-CoV-2, COVID-19, senescence, NK cells, CD8^+^ T cells

## Abstract

Unvaccinated COVID-19 patients display a large spectrum of symptoms, ranging from asymptomatic to severe symptoms, the latter even causing death. Distinct Natural killer (NK) and CD4^+^ and CD8^+^ T cells immune responses are generated in COVID-19 patients. However, the phenotype and functional characteristics of NK cells and T-cells associated with COVID-19 pathogenesis versus protection remain to be elucidated. In this study, we compared the phenotype and function of NK cells SARS-CoV-2-specific CD4^+^ and CD8^+^ T cells in unvaccinated symptomatic (SYMP) and unvaccinated asymptomatic (ASYMP) COVID-19 patients. The expression of senescent CD57 marker, CD45RA/CCR7differentiation status, exhaustion PD-1 marker, activation of HLA-DR, and CD38 markers were assessed on NK and T cells from SARS-CoV-2 positive SYMP patients, ASYMP patients, and Healthy Donors (HD) using multicolor flow cytometry. We detected significant increases in the expression levels of both exhaustion and senescence markers on NK and T cells from SYMP patients compared to ASYMP patients and HD controls. In SYMP COVID-19 patients, the T cell compartment displays several alterations involving naive, central memory, effector memory, and terminally differentiated T cells. The senescence CD57 marker was highly expressed on CD8^+^ T_EM_ cells and CD8^+^ T_EMRA_ cells. Moreover, we detected significant increases in the levels of proinflammatory TNF-α, IFN-γ, IL-6, IL-8, and IL-17 cytokines from SYMP COVID-19 patients, compared to ASYMP COVID-19 patients and HD controls. The findings suggest exhaustion and senescence in both NK and T cell compartment is associated with severe disease in critically ill COVID-19 patients.

**IMPORTANCE:** Unvaccinated COVID-19 patients display a large spectrum of symptoms, ranging from asymptomatic to severe symptoms, the latter even causing death. Distinct Natural killer (NK) and CD4^+^ and CD8^+^ T cells immune responses are generated in COVID-19 patients. In this study, we detected significant increases in the expression levels of both exhaustion and senescence markers on NK and T cells from unvaccinated symptomatic (SYMP) compared to unvaccinated asymptomatic (ASYMP) COVID-19 patients. Moreover, we detected significant increases in the levels of proinflammatory TNF-α, IFN-γ, IL-6, IL-8, and IL-17 cytokines from SYMP COVID-19 patients, compared to ASYMP COVID-19 patients. The findings suggest exhaustion and senescence in both NK and T cell compartment is associated with severe disease in critically ill COVID-19 patients.

**TWEET:** Significant exhaustion and senescence in both NK and T cells were detected in unvaccinated symptomatic COVID-19 patients, suggesting a weakness in both innate and adaptive immune systems leads to severe disease in critically ill COVID-19 patients.

## INTRODUCTION

Severe acute respiratory syndrome coronavirus 2 (SARS-CoV-2) is a β-Coronavirus that was first detected in 2019 in Wuhan, China. In the ensuing months, it has been transmitted worldwide. As of July 2022, more than 568 million people have contracted coronavirus disease (COVID-19), the pandemic that has killed approximately 6.38 million people globally (1). COVID-19, caused by SARS CoV-2, has a wide range of clinical manifestations, ranging from asymptomatic to severe symptomatic disease (2). Therefore, understanding the clinical and immunological characteristics of unvaccinated ASYMP and SYMP COVID-19 patients holds significance in elucidating the immunopathogenesis of COVID-19 and informing the development of effective immune treatments. Within 2-14 days after SARS-CoV-2 exposure, newly infected individuals may develop fever, fatigue, myalgia, and respiratory symptoms, including cough and shortness of breath (3, 4). While the majority (80-85%) of newly infected individuals are asymptomatic (i.e., patients who remain symptomless despite being SARS-CoV-2-positive), a minority of individuals are symptomatic, especially the elderly and those with compromised health, that develop severe pulmonary inflammatory disease and may need a rapid medical intervention to prevent acute respiratory distress syndrome and death (5-10).

Innate and adaptive immune responses are of great significance for the control of viral infections. NK cells exert the primary control during acute viral infection, but CD4^+^ and cytotoxic CD8^+^ T lymphocytes (CTLs) are critical for the long-term surveillance (11). Recently, De Biasi et al. reported an increase in the CD57 expression on CD8^+^ T cells (12, 13). CD57 is a key marker of *in vitro* replicative senescence and is associated with prolonged chronic infection (14). Immunosenescence includes a shift towards less functional T cells in the immune system (15). However, CD57 expression is reported to be a marker of mature NK cells. The phenotypes and differentiation status associated with replicative senescent T lymphocytes are not well-defined. Like T-cells, NK cell expression of CD57 could be considered as a marker of terminal differentiation (16). Furthermore, the expression of CD57 aids in identifying the final stages of peripheral NK cell maturation, and the expression of CD57 increases with age and chronic infections (16).

Reports show that repeated T cell activation is associated with terminally differentiated cells and the corresponding upregulation of CD57 (13, 17). It is observed that shortened telomeres are features of senescent cells, and replicative senescence results in a low proliferative capacity of the cells, eventually leading to an inability to eradicate infection (18).

Understanding the spectrum of innate and adaptive immune responses against SARS-CoV2, disease severity, and cellular immunosenescence in SARS-CoV-2 infected symptomatic versus asymptomatic individuals can ultimately inform the identification of new therapeutic targets. To attain this goal, we phenotypically and functionally characterized the senescence markers (CD57), differentiation status (CD45RA/CCR7), exhaustion marker (PD-1), and activation marker (HLA-DR and CD38) from patients with SARS-CoV-2 positive symptomatic and asymptomatic patients, and Healthy controls using multicolor flow cytometry.

In this report, we show 1) a decreased CD56^bright^ NK cell population and higher frequency of mature/terminally differentiated NK cells (CD57^+^) in SYMP patients; 2) the activation status, senescence, and exhaustion profile were significantly increased in COVID-19 SYMP individuals; 3) SARS-CoV-2 specific senescent T cells with an effector memory phenotype (CD57^+^ CD8^+^ T_EM_ and CD57^+^ CD8^+^ T_EMRA_ cells) was detected in COVID-19 SYMP individuals; 4) COVID-19 patients displayed increased cytokine storm detectable in the plasma samples.

Our findings demonstrate that increased T cell exhaustion and senescence markers in unvaccinated ASYMP COVID-19 patients compared to unvaccinated ASYMP COVID-19 patients and Healthy Controls. Furthermore, T cell senescence markers were highly expressed on CD8^+^ T_EM_ and CD8^+^ T_EMRA_ cells than on T_NAIVE_ and T_CM_ cells. These results suggest that the upregulation of exhaustion and senescence pathways during symptomatic COVID-19 may affect both NK and T cell compartments, leading to inefficient clearance of SARS-CoV-2 infection and severe disease.

## MATERIALS & METHODS

### Human study population

All clinical investigations in this study were conducted according to the Declaration of Helsinki principles. All subjects were enrolled at the University of California, Irvine, under approved Institutional Review Board-approved protocols (IRB#-2020-5779). Written informed consent was received from all participants before inclusion. Twenty COVID-19 patients (Asymptomatic and Symptomatic) and ten unexposed Healthy individuals, who had never been exposed to SARS-CoV-2 or COVID-19 patients, were enrolled in this study (**Table 1**). Thirty percent were Caucasian, and 70% were non-Caucasian. Forty-four percent were females, and 60% were males with an age range of 21-67 years old (median 39). None of the symptomatic patients were on anti-viral or anti-inflammatory drug treatments during blood sample collections.

**Table I.**
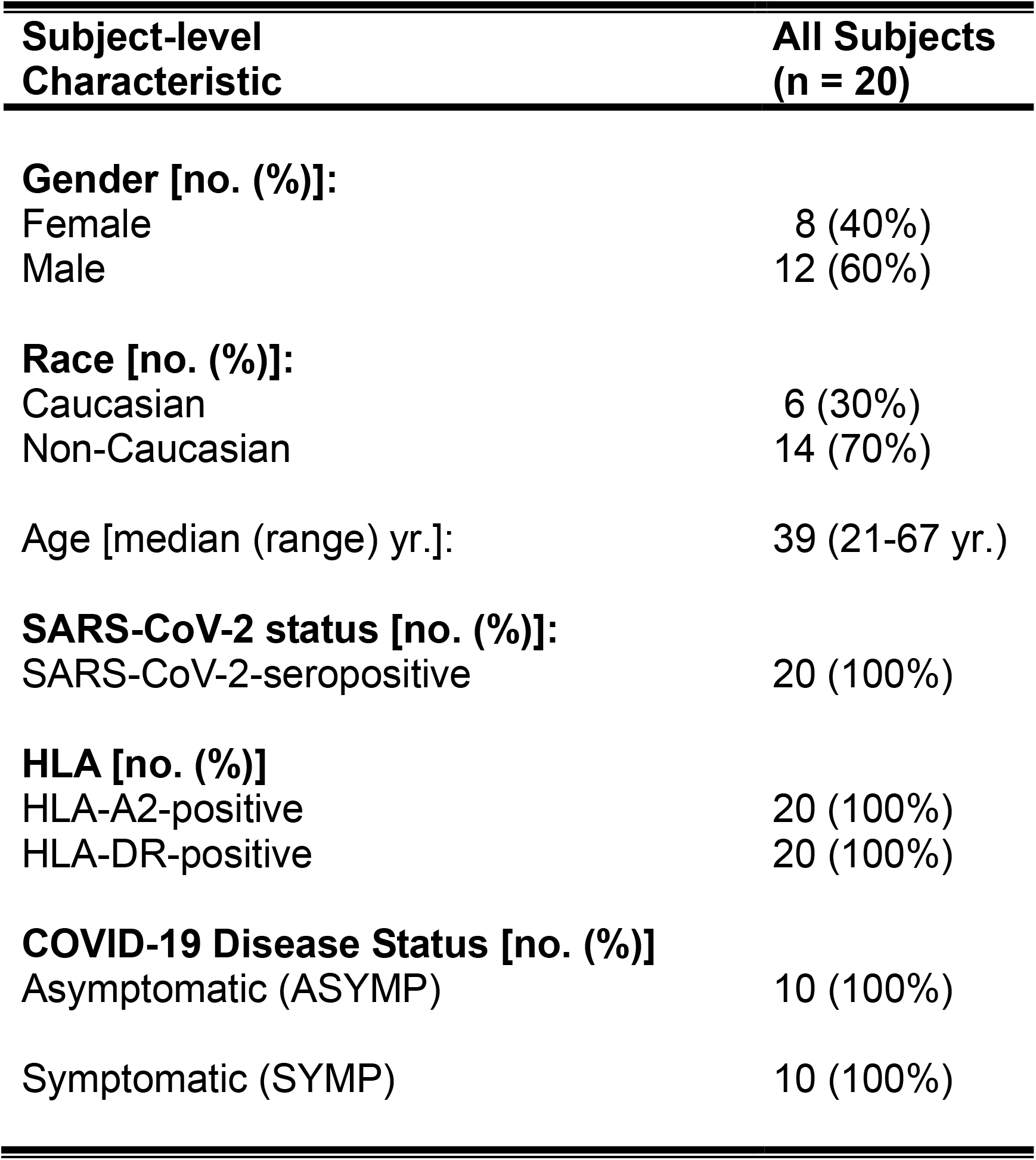
Cohorts of HLA-A2/HLA-DR positive, SARS-CoV-2 seropositive Symptomatic and Asymptomatic individuals enrolled in this study

| **Subject-level Characteristic** | **All Subjects (n = 20)** |
| --- | --- |
| **Gender [no. (%)]:** |  |
| Female | 8 (40%) |
| Male | 12 (60%) |
| **Race [no. (%)]:** |  |
| Caucasian | 6 (30%) |
| Non-Caucasian | 14 (70%) |
| Age [median (range) yr.]: | 39 (21-67 yr.) |
| **SARS-CoV-2 status [no. (%)]:** |  |
| SARS-CoV-2-seropositive | 20 (100%) |
| **HLA [no. (%)]** |  |
| HLA-A2-positive | 20 (100%) |
| HLA-DR-positive | 20 (100%) |
| **COVID-19 Disease Status [no. (%)]** |  |
| Asymptomatic (ASYMP) | 10 (100%) |
| Symptomatic (SYMP) | 10 (100%) |

Detailed clinical and demographic characteristics of the symptomatic versus asymptomatic COVID-19 patients and the unexposed Healthy individuals concerning age, gender, HLA-A*02:01, and HLA-DR distribution, COVID-19 disease severity, comorbidity, and biochemical parameters are detailed in **Table 1**.

### HLA-A2 typing

The HLA-A2 status was confirmed by PBMCs staining with 2 μL of anti-HLA-A2 mAb (clone BB7.2) (BD Pharmingen, Franklin Lakes, NJ), at 4°C for 30 minutes. The cells were washed and analyzed by flow cytometry using an LSRII (Becton Dickinson, Franklin Lakes, NJ). The acquired data were analyzed with FlowJo software (BD Biosciences, San Jose, CA).

### Tetramer/ peptide staining

Fresh PBMCs were analyzed for the frequency of CD8^+^ T cells recognizing the SARS-CoV-2 peptide/tetramer complexes, as we previously described in (19). The cells were incubated with SARS-CoV-2 peptide/tetramer complex for 30–45 min at 37°C. The cell preparations were then washed with FACS buffer and stained with FITC-conjugated anti-human CD8 mAb (BD Pharmingen). Finally, the cells were washed and fixed with 1% paraformaldehyde in PBS and subsequently acquired on a BD LSRII. Data were analyzed using FlowJo version 9.5.6 (Tree Star).

### Human peripheral blood mononuclear cells (PBMC) isolation

SARS-COV-2 positive individuals were recruited at the UC Irvine Medical Center. Between 40 -50 mL of blood was drawn into yellow-top Vacutainer^®^ Tubes (Becton Dickinson). The plasma samples were isolated and stored at -80° C for the detection of various cytokines using Luminex. PBMCs were isolated by gradient centrifugation using a leukocyte separation medium (Life Sciences, Tewksbury, MA). The cells were then washed in PBS, and re-suspended in a complete culture medium consisting of RPMI1640, 10% FBS (Bio-Products, Woodland, CA) supplemented with 1x penicillin/streptomycin/L-glutamine, 1x sodium pyruvate, 1x non-essential amino acids, and 50 μM of 2-mercaptoethanol (Life Technologies, Rockville, MD). For future testing, freshly isolated PBMCs were also cryopreserved in 90% FCS and 10% DMSO in liquid nitrogen.

### Human T cells flow cytometry assays

The following anti-human antibodies were used for the flow cytometry assays: CD3 Percp, CD8 APC-Cy7, CD57 PE-Cy7, PD-1 A647, CD45RA FITC, CCR7 BV786, HLA-DR BUV385, CD38 A700, CD56 APC (BioLegend, San Diego, CA). For surface staining, mAbs against cell markers were added to a total of 1 × 10^6^ cells in 1X PBS containing 1% FBS and 0.1% sodium azide (FACS buffer) for 45 minutes at 4°C. After washing with FACS buffer, cells were permeabilized for 20 minutes on ice using the Cytofix/Cytoperm Kit (BD Biosciences) and then washed twice with Perm/Wash Buffer (BD Biosciences). Intracellular cytokine mAbs were then added to the cells and incubated for 45 minutes on ice in the dark. Finally, cells were washed with Perm/Wash and FACS Buffer and fixed in PBS containing 2% paraformaldehyde (Sigma-Aldrich, St. Louis, MO). For each sample, 100,000 total events were acquired on the BD LSRII. Ab capture beads (BD Biosciences) were used as individual compensation tubes for each fluorophore in the experiment. We used fluorescence minus controls for each fluorophore to define positive and negative populations when initially developing staining protocols. In addition, we further optimized gating by examining known negative cell populations for background expression levels. The gating strategy was similar to that used in our previous work (20). Briefly, we gated on single cells, dump cells, viable cells (Aqua Blue), lymphocytes, CD3^+^ cells, and human epitope-specific CD8^+^ T cells using HSV-specific tetramers. Data analysis was performed using FlowJo version 9.9.4 (TreeStar, Ashland, OR). Statistical analyses were done using GraphPad Prism version 5 (La Jolla, CA).

### Statistical analyses

Data for each assay were compared by analysis of variance (ANOVA) and Student’s *t*-test using GraphPad Prism version 5.03. ANOVA and multiple comparison procedures identified differences between the groups, as we previously described in (21). Data are expressed as the mean + SD. Results were considered statistically significant at *p* < 0.05.

## RESULTS

### 1. Composition of NK cell subsets in COVID-19 SYMP patients show a decreased CD56^bright^ NK cell population and higher frequency of mature/terminally differentiated NK cells (CD57^+^) compared to Healthy individuals

NK cells are a subset of innate immune lymphocytes composing 5% to 20% of PBMCs in humans and play an important role in the defense against viral infections. These cells are reduced in numbers but less consistently than T cells, particularly in severely sick patients. Therefore, we first investigated the phenotypic status of NK cells in SARS-CoV-2 positive asymptomatic (ASYMP) and symptomatic (SYMP) patients and Healthy Controls. The characteristics of the SYMP, ASYMP and Healthy control study populations used in this study, concerning age, sex, HLA-A*02:01 frequency distribution, SARS-CoV-2 positivity, and status of COVID-19 disease are presented in **Table 1**. These SARS-CoV-2 positive individuals were divided into two groups: 1) HLA-A*02:01–positive SARS-CoV-2–infected ASYMP individuals, with no detectable levels of any clinical COVID-19 disease; and 2) HLA-A*02:01–positive SARS-CoV-2– infected SYMP individuals with a well-documented COVID-19 clinical disease.

We analyzed the NK cell population following a gating strategy as shown in **Fig. 1A**. The NK cell population was further categorized into CD56^dim^ and CD56^bright^ cells, and their relative frequency was evaluated in ASYMP, SYMP and Healthy individuals. Analysis of NK cell phenotype showed no difference in mature CD56^dim^ subset in SARS-CoV-2 positive ASYMP and SYMP patients and Healthy controls (**Fig. 1B**, *top panel*). However, SYMP patients significantly reduced immature CD56^bright^ NK cells (**Fig. 1B**, *bottom panel*; *P* = 0.02) compared to Healthy controls.

**Figure 1:**
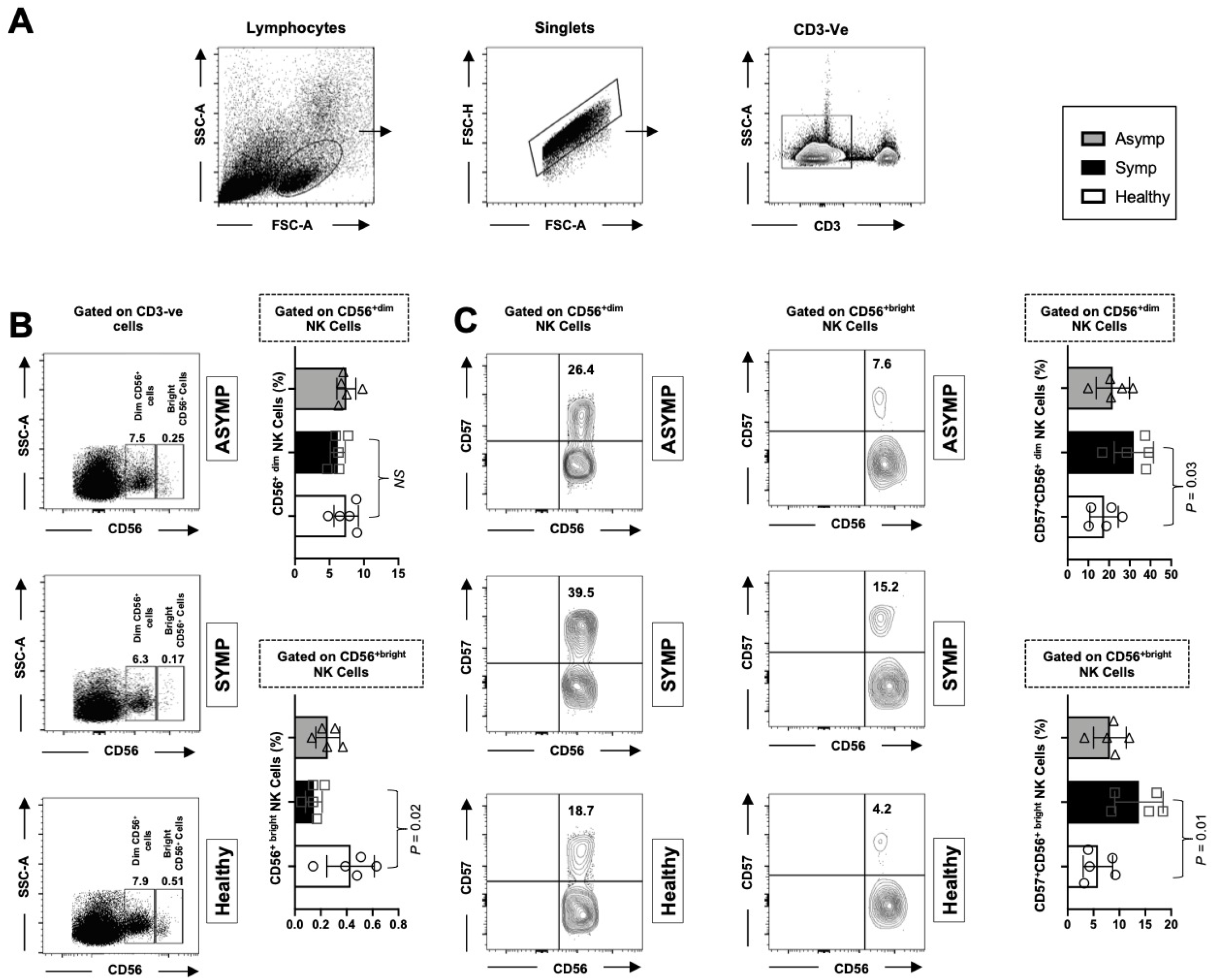
Composition of NK cell subsets in COVID-19 SYMP individuals shows a decreased CD56^bright^ NK cell population and a higher frequency of mature/terminally differentiated NK cells (CD57^+^) than in Healthy individuals.A gating strategy for defining of NK cell population is shown using FACS. (**A**) Using forward scatter (FSC) and side scatters (SSC), the lymphocyte populations were gated. Singlets were gated after gating the lymphocytes population, and NK cells were defined as CD3^-^ CD56^+^ cells. The NK cell population was further categorized into CD56^dim^ and CD56^bright^ cells, and their relative frequency of senescence (CD57^+^) was evaluated. (**B**) Representative FACS data of the frequencies of CD56^dim^ NK cells and CD56^bright^ NK cells detected in PBMCs from COVID-19 ASYMP individuals SYMP individuals and Healthy controls (*left panel*). Average frequencies of PBMC-derived CD56^dim^ NK cells (*top*) and CD56^bright^ NK cells (*bottom*) were detected from ASYMP, SYMP and Healthy individuals (*right panel*). (**C**) Representative FACS data of the frequencies of CD57 gated on CD56^dim^ NK cells and the frequencies of CD57 gated on CD56^bright^ NK cells detected in PBMCs from COVID-19 ASYMP individual, SYMP individual, and Healthy control (*left panel*). Average frequencies of PBMCs-derived CD56^dim^ NK cells (*top*) and CD56^bright^ NK cells (*bottom*) were detected from ASYMP, SYMP and Healthy individuals (*right panel*). The results are representative of two independent experiments on each individual. The indicated *P* values, calculated using an unpaired t-test, show statistical significance between SYMP and Healthy individuals.

CD57 expression on NK cells defines a mature phenotype, and their expression of CD57 could also be considered a marker of terminal differentiation, although not associated with senescence in this population. It is highly expressed on CD56^dim^ cells, representing mature NK cells, whereas less than 1% of CD56^bright^ NK cells, considered immature, also express CD57 (**Fig. 1C**). There was a significant increase in the mature (CD56^dim^ /CD57^+^) and immature subset (CD56^bright^ /CD57^+^) (**Fig. 1C**, *top* and *bottom panel*; *P* = 0.03 and *P* = 0.01 respectively) in SYMP patients with COVID-19 compared with Healthy controls.

Collectively, these data indicate different states of maturation within the CD56^dim^ and CD56^bright^ NK-cell subset and its correlation with COVID-19.

### 2. The activation status, senescence, and exhaustion profile were significantly increased in COVID-19 SYMP individuals compared to Healthy individuals within CD4^+^ T cells

CD4^+^ T cells in COVID-19 are activated as characterized by the expression of cellular markers like HLA-DR and CD38. Therefore, we next evaluated the degree of CD4^+^ T cell activation in COVID-19 positive ASYMP and SYMP patients and Healthy Controls. Within the CD4^+^ population, we analyzed markers commonly related to T cell activation (HLA-DR and CD38). The gating strategy used to analyze markers related to activation status, senescence, and exhaustion together within CD4^+^ T cells is demonstrated in **Fig. 2A**. The expression of CD57 correlates with senescence in human CD4^+^ and CD8^+^ T cells. Therefore, we compared the frequency of CD57^+^ on total CD4^+^ T cells in COVID-19 ASYMP, SYMP and Healthy individuals. PBMC-derived CD57^+^CD4^**+**^ T cells detected from COVID-19 SYMP individuals showed an increased frequency compared to Healthy individuals (**Fig. 2B;** *P* = 0.03).

**Figure 2:**
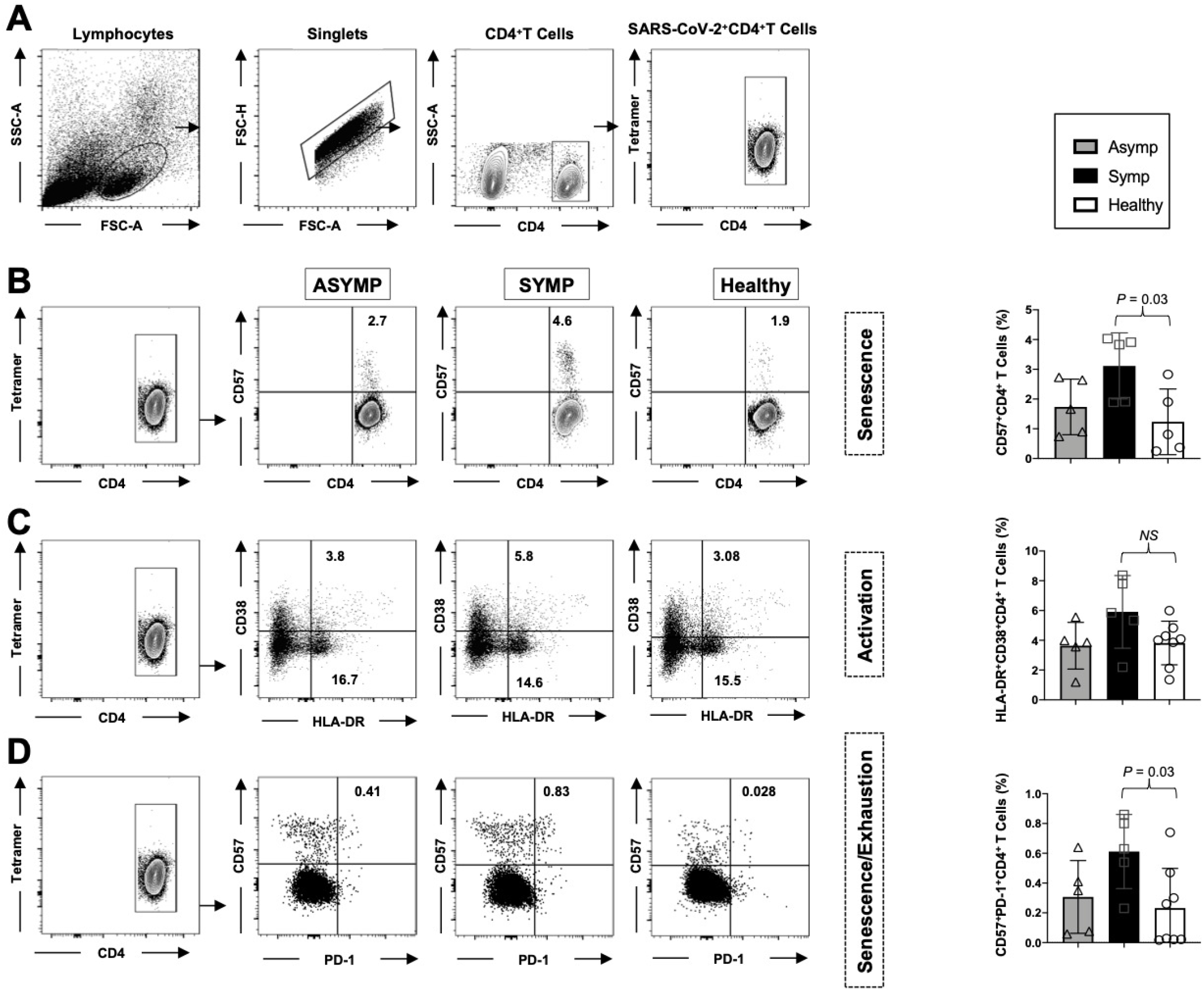
The activation status, senescence, and exhaustion profile were significantly increased in COVID-19 SYMP individuals compared to Healthy individuals within CD4^+^ T cells. Expression of CD38 and HLA-DR was detected to analyze the activation status of CD4^+^ T cells. Expression of CD57 and PD-1 was detected to analyze the senescence/exhaustion status of CD4^+^ T cells. (**A**) The gating strategy was used to analyze markers related to activation status, senescence, and exhaustion together within CD4^+^ T cells. Activated cells are CD38^+^HLA-DR^+^ ; exhausted/senescent are PD1^+^CD57^+^. (**B**) Representative FACS data of the frequencies of CD57^+^CD4^+^ T cells detected in PBMCs from COVID-19 ASYMP, SYMP and Healthy individuals (*left panel*). Average frequencies of PBMC-derived CD57^+^CD4^+^ T cells were detected from ASYMP, SYMP and Healthy individuals (*right panel*). (**C**) Representative FACS data of the frequencies of HLA-DR^+^CD38^+^CD4^+^ T cells detected in PBMCs from ASYMP individual, SYMP individual and Healthy control (*left panel*). Average frequencies of PBMC-derived HLA-DR^+^CD38^+^CD4^+^ T cells were detected from ASYMP SYMP and Healthy individuals (*right panel*). (**D**) Representative FACS data of the frequencies of CD57^+^PD-1^+^CD4^+^ T cells detected in PBMCs from COVID-19 ASYMP, SYMP and Healthy individuals (*left panel*). Average frequencies of PBMCs-derived CD57^**+**^ PD-1^+^CD4^**+**^ T cells were detected from ASYMP, SYMP and Healthy individuals (*right panel*). The results are representative of two independent experiments on each individual. The indicated *P* values, calculated using an unpaired t-test, show statistical significance between SYMP and Healthy individuals.

The levels of CD4^+^ T-cell activation were also evaluated in SARS-CoV-2 infected ASYMP, SYMP patients, and healthy controls. We found an increasing trend reflected by higher proportions of HLA-DR^+^CD38^+^CD4^+^ T cells in SYMP patients compared to healthy controls, however with no statistical significance (**Fig. 2C**).

Furthermore, we evaluated the expression of the senescence/exhaustion molecules on CD4^+^ T cells by analyzing markers CD57 and PD-1 (**Fig. 2D)**. The proportion of CD57^+^PD-1^+^CD4^+^ T cells was significantly higher in COVID-19 positive SYMP patients than in healthy controls and ASYMP patients.

However, no statistical difference was detected in the proportion of CD57^+^PD-1^+^CD4^+^ T cells in ASYMP and SYMP patients (**Fig. 2D;** *P* = 0.03).

Taken together, findings from COVID-19 positive SYMP individuals indicate the involvement of activated CD4^+^ T cells and T cell exhaustion/senescence in the immunopathogenesis of SARS-CoV2 infection.

### 3. Frequent SARS-CoV-2 specific senescent CD4^+^ T cells with an effector memory phenotype (CD57^+^CD4^+^ T_EM_ and CD57^+^CD4^+^ T_EMRA_ cells) detected in COVID-19 SYMP individuals compared to Healthy individuals

SARS-CoV-2 specific memory CD4^+^ T cells were also categorized into three major phenotypically distinct effector memory (TEM), central memory (TCM) and a subset of effector memory T cells re-expresses CD45RA subpopulations termed as TEMRA. Moreover, we studied the expression levels of CD57 on the memory CD4^+^ T cell subpopulations at various stages of differentiation: central memory T cells, (CD45RA^low^ CCR7^high^ CD4^+^ T_CM_ cells); effector memory T cells, (CD45RA^low^ CCR7^low^ CD4^+^ T_EM_ cells) and TEMRA T cells (CD45RA^high^ CCR7^low^ CD4^+^ T_EMRA_ cells). In the peripheral blood of HLA-A*02:01 positive, SARS-CoV-2 positive ASYMP, SYMP and Healthy individuals, we compared the CD57 expression in CD4^+^ T cells and divided them into T_NAIVE_, T_CM_, T_EM_, and T_EMRA_ phenotypes (**Fig. 3A**). Similar percentages of CD57^+^ CD45RA^low^ CCR7^high^ CD4^+^ T_CM_ cells were detected in ASYMP, SYMP and Healthy individuals (**Fig. 3B;** *left* and *right* panel). There was an increase in the CD57^+^ CD45RA^low^ CCR7^low^ CD4^+^ T_EM_ cells in SYMP individuals compared to Healthy controls (**Fig. 3C;** *left* and *right* panel (*P*=0.02)). Significantly higher percentages of CD57^+^ CD45RA^high^ CCR7^low^ CD4^+^ T_EMRA_ cells (**Fig. 3D;** *left* and *right* panel) were detected in SYMP individuals compared to Healthy individuals (*P* = 0.01). Altogether, the phenotypic properties of SARS-CoV-2 specific memory CD4^+^ T cells revealed a clear dichotomy in memory CD4^+^ T cell sub-populations in SYMP versus Healthy individuals. SYMP individuals appeared to develop frequent effector memory CD57^+^CD4^+^ T_EMRA_ and CD57^+^CD4^+^ T_EM_ cells compared to Healthy and ASYMP individuals. By maintaining high frequencies of the SARS-CoV-2-specific CD57^+^CD4^+^ T_EMRA_ cells and CD57^+^CD4^+^ T_EM_ cells, the SYMP individuals may not be protected against infection and/or COVID-19 disease.

**Figure 3:**
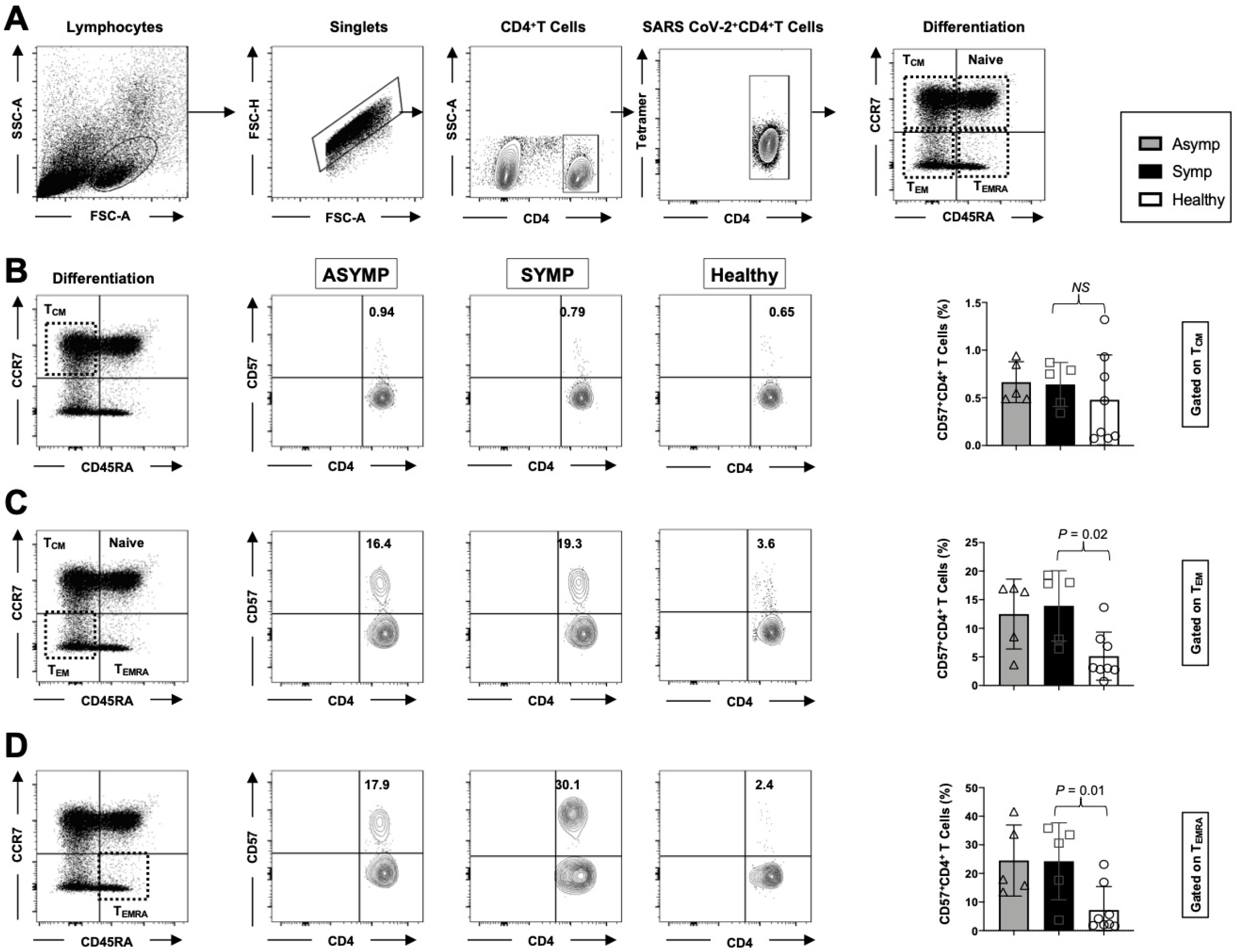
Frequent SARS-CoV-2 specific senescent CD4^+^ T cells with an effector memory phenotype (CD57^+^CD4^+^ T_EM_ and CD57^+^CD4^+^ T_EMRA_ cells) detected in COVID-19 SYMP individuals compared to Healthy individuals. The phenotype of CD4^+^ T cells and the gating strategy shown in **Fig. 2A** were analyzed in T_NAIVE_, T_CM_, T_EMRA_, and T_EM_ phenotypes in PBMCs from COVID-19 ASYMP SYMP and Healthy individuals. Representative FACS data (*left panel*) and the frequencies of CD57 (*right panel*) (**B**) gated on CD45RA^low^ CCR7^high^ CD4^+^ T_CM_ cells, (**C**) gated on CD45RA^low^ CCR7^low^ CD4^+^ T_EM_ cells, and (**D**) and CD45RA^high^ CCR7^low^ CD4^+^ T_EMRA_ cells detected in COVID-19 ASYMP individual, SYMP individual and Healthy individual. The results are representative of two independent experiments on each individual. The indicated *P* values, calculated using an unpaired t-test, show statistical significance between SYMP and Healthy individuals.

### 4. Frequent SARS-COV-2 S_1220-1228_ and S_958-966_ epitope-specific CD57^+^ CD8^+^ T cells detected in COVID-19 SYMP individuals compared to ASYMP and Healthy individuals

As described earlier SARS-CoV-2 positive individuals were segregated into two groups: 1) HLA-A*02:01– positive SARS-CoV-2–infected ASYMP individuals, and 2) HLA-A*02:01–positive SARS-CoV-2– infected SYMP individuals with a well-documented COVID-19 clinical disease. We next compared the frequency of CD57^+^ on total CD8^+^ T cells in HLA-A*02:01 positive COVID-19 ASYMP, SYMP and Healthy individuals. We have used a gating strategy to analyze markers related to senescence (CD57^+^) gated within CD8^+^ T cells from COVID-19 ASYMP, SYMP and Healthy individuals. Average frequencies of PBMC-derived CD57^+^ CD8^**+**^ T cells detected from COVID-19 SYMP individuals showed an increased frequency compared to Healthy individuals (**Fig. 4A**, *P* = 0.03). We then compared the frequency of SARS-CoV-2 peptide/tetramer complex specific CD8^+^ T cells. The representative dot plots in **Fig. 4B** indicate an increased frequency of CD57^+^ CD8^+^ T cells, specific to S1220-1228 epitope in COVID-19 SYMP individuals compared to ASYMP Healthy individuals (*P* = 0.01). Similarly, **Fig. 4C** depicts the high frequencies of CD57^+^ CD8^+^ T cells detected in COVID-19 SYMP individuals against another peptide/tetramer complex S_958-966_ epitope (*P* = 0.02). Altogether, these results indicate that SYMP individuals develop frequent SARS-CoV-2-specific CD57^+^ CD8^+^ T cells compared to ASYMP and Healthy individuals.

**Figure 4:**
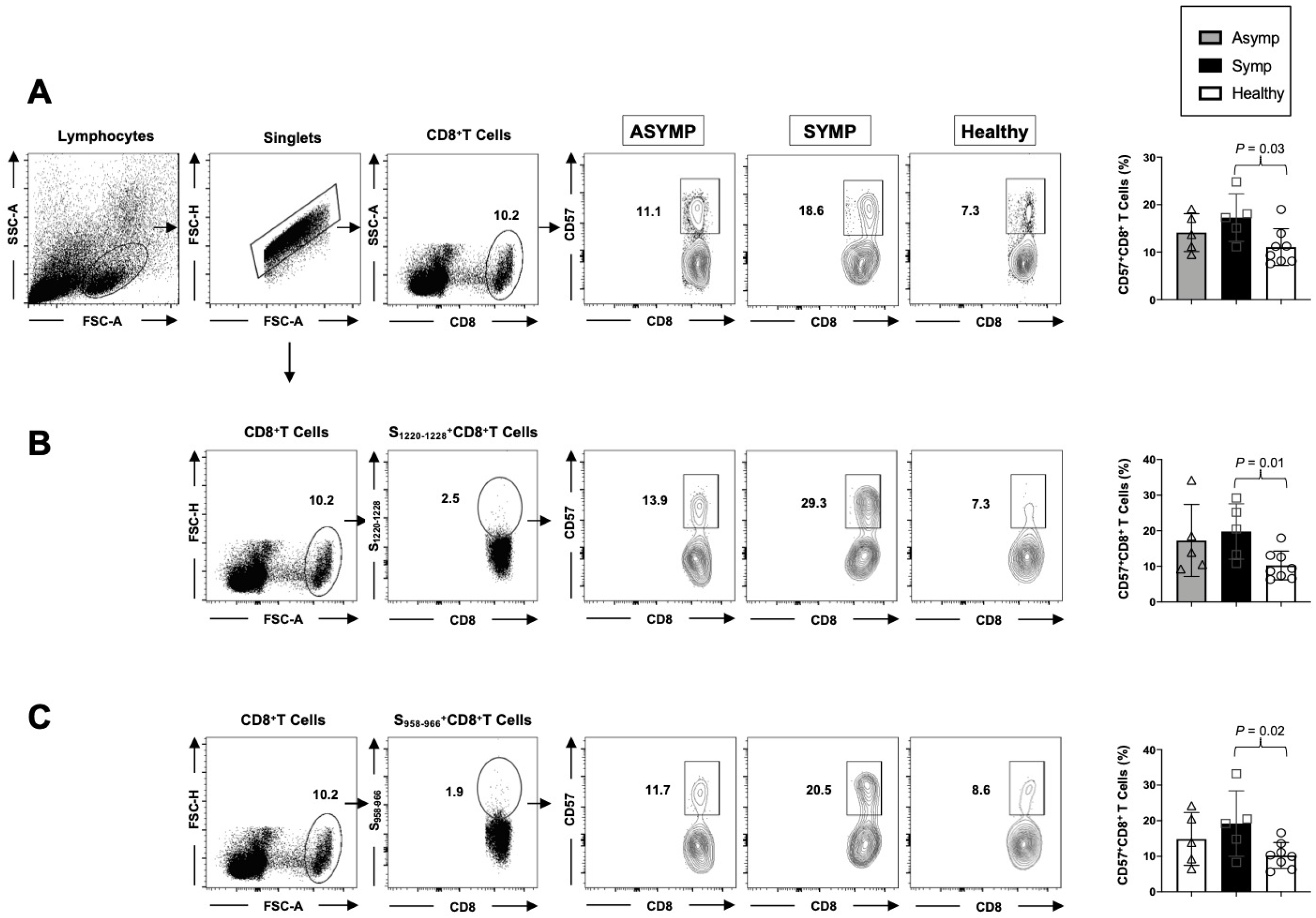
Frequent SARS-COV-2 S_1220-1228_ and S_958-966_ epitope-specific CD57^+^ CD8^+^ T cells detected in COVID-19 SYMP individuals compared to ASYMP and Healthy individuals. The frequency of CD57^+^ on total CD8^+^ T cells and SARS-CoV-2 peptide/tetramer complex specific CD8^+^ T cells were analyzed in HLA-A*02:01 positive COVID-19 ASYMP, SYMP and Healthy individuals. (**A**) Gating strategy used to analyze markers related to senescence gated within CD8^+^ T cells from COVID-19 ASYMP, SYMP and Healthy individuals (*left panel*), and average frequencies of PBMC-derived CD8^**+**^ T cells detected from COVID-19 ASYMP, SYMP and Healthy individuals (*right panel*). (**B**) Representative FACS data of the frequencies of CD57^+^ CD8^+^ T cells, specific to S_1220-1228_ epitope, detected in PBMCs from HLA-A*02:01 positive COVID-19 ASYMP, SYMP and Healthy individuals (*left panel*). Average frequencies of PBMC-derived CD8^**+**^ T cells, specific to S_1220-1228_ epitope, were detected from COVID-19 ASYMP, SYMP and Healthy individuals (*right panel*). (**C**) Representative FACS data of the frequencies of CD57^+^ CD8^+^ T cells, specific to S_958-966_ epitope, detected in PBMCs from HLA-A*02:01 positive COVID-19 ASYMP, SYMP and Healthy individuals (*left panel*). Average frequencies of PBMC-derived CD57^**+**^ CD8^**+**^ T cells, specific to S_958-966_ epitope, were detected from ASYMP, SYMP and Healthy individuals (*right panel*). The results are representative of two independent experiments on each individual. The indicated *P* values, calculated using an unpaired t-test, show statistical significance between SYMP and Healthy individuals.

### 5. Frequent SARS-CoV-2 S_1220-1228_ epitope-specific senescent CD8^+^ T cells with an effector memory phenotype (CD57^+^ CD8^+^ T_EM_ and CD57^+^ CD8^+^ T_EMRA_ cells) detected in COVID-19 SYMP individuals compared to Healthy individuals

Similar to the CD4^+^ T cell memory response, SARS-CoV-2 specific memory CD8^+^ T cells are also categorized into three major phenotypically distinct effector memory (TEM), central memory (TCM) and a subset of effector memory T cells reexpresses CD45RA subpopulations termed as TEMRA. Furthermore, we examined the expression levels of CD57 on the memory CD8^+^ T cell subpopulations at various stages of differentiation: central memory T cells, (CD45RA^low^ CCR7^high^ CD8^+^ T_CM_ cells); effector memory T cells, (CD45RA^low^ CCR7^low^ CD8^+^ T_EM_ cells) and TEMRA T cells (CD45RA^high^ CCR7^low^ CD8^+^ T_EMRA_ cells). In the peripheral blood of HLA-A*02:01 positive, SARS-CoV-2 positive ASYMP, SYMP and Healthy individuals, we compared the CD57 expression in CD8^+^ T cells specific to S_1220-1228_ epitope and divided them into T_NAIVE_, T_CM_, T_EM,_ and T_EMRA_ phenotypes (**Fig. 5A**). Similar percentages of CD57^+^ CD45RA^low^ CCR7^high^ CD8^+^ T_CM_ cells were detected in ASYMP, SYMP and Healthy individuals (**Fig. 5B;** *left* and *right* panel). There was a significant increase in the CD57^+^ CD45RA^low^ CCR7^low^ CD8^+^ T_EM_ cells in SYMP individuals compared to Healthy controls (**Fig. 5C;** *left* and *right* panel). Significantly higher percentages of CD57^+^ CD45RA^high^ CCR7^low^ CD8^+^ T_EMRA_ cells (**Fig. 5D;** *left* and *right* panel) were detected in SYMP individuals compared to Healthy individuals (*P* = 0.004). Altogether, the phenotypic properties of SARS-CoV-2 specific S_1220-1228_ epitope epitope-specific memory CD8^+^ T cells revealed a clear dichotomy in memory CD8^+^ T cell sub-populations in SYMP versus Healthy individuals. SYMP individuals appeared to develop frequent SARS-CoV-2 specific effector memory CD57^+^ CD8^+^ T_EMRA_ and CD57^+^ CD8^+^ T_EM_ cells compared to Healthy and ASYMP individuals. By maintaining high frequencies of the SARS-CoV-2-specific CD57^+^ CD8^+^ T_EMRA_ cells and CD57^+^ CD8^+^ T_EM_ cells, the SYMP individuals may not be protected against infection and/or COVID-19 disease.

**Figure 5:**
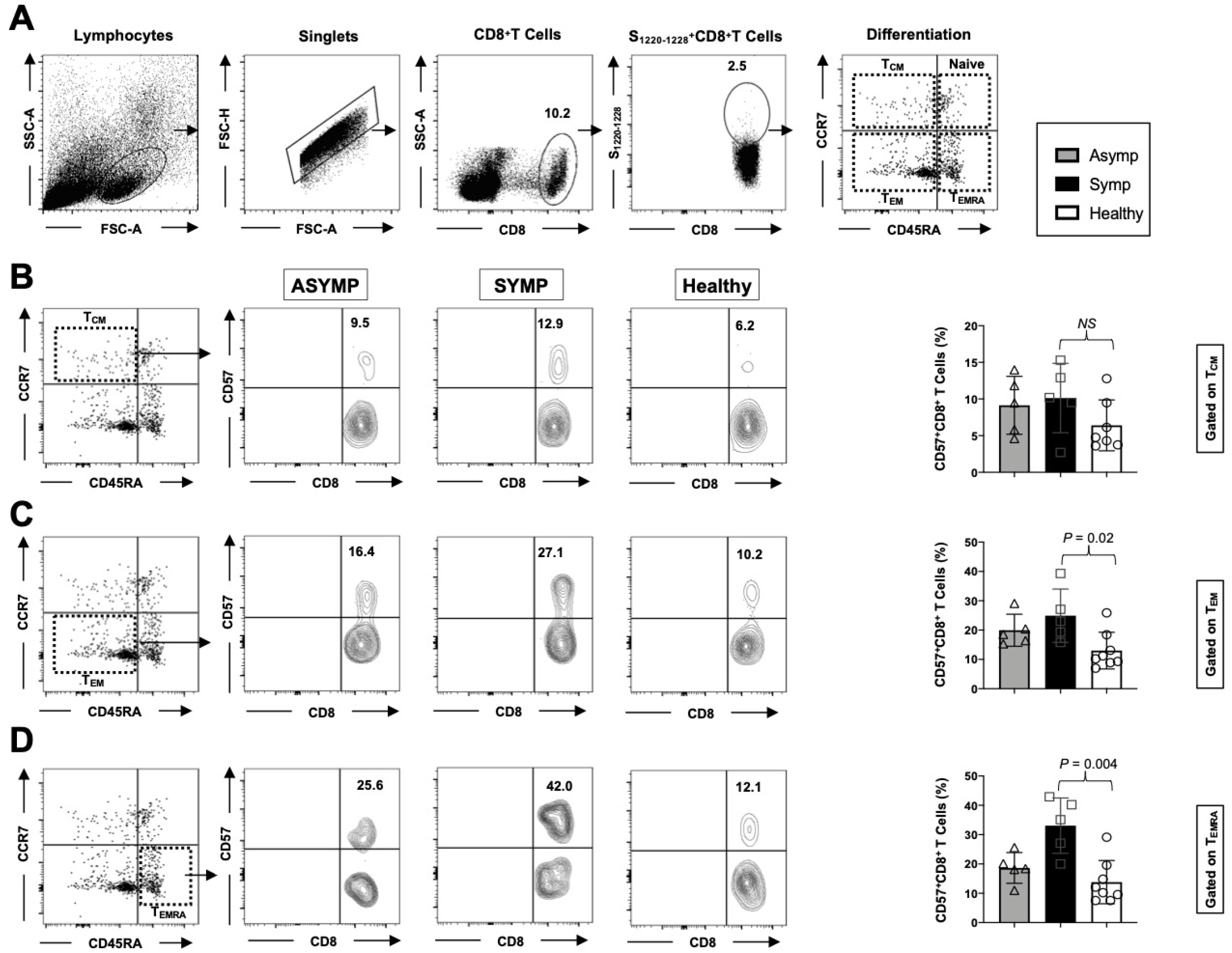
Frequent SARS-CoV-2 S_1220-1228_ epitope-specific senescent CD8^+^ T cells with an effector memory phenotype (CD57^+^ CD8^+^ T_EM_ and CD57^+^ CD8^+^ T_EMRA_ cells) detected in COVID-19 SYMP individuals compared to Healthy individuals. The phenotype of CD8^+^ T cells specific to S_1220-228_ peptide/tetramer shown in **Fig. 5A** was analyzed in terms of T_NAIVE_, T_CM_, T_EMRA,_ and T_EM_ phenotypes in PBMCs from HLA-A*02:01 positive COVID-19 ASYMP, SYMP and Healthy individuals. Representative FACS data (*left panel*) and the frequencies of CD57 (*right panel*) gated on CD45RA^low^ CCR7^high^ CD8^+^ T_CM_ cells (**B**), gated on CD45RA^low^ CCR7^low^ CD8^+^ T_EM_ cells (**C**), and CD45RA^high^ CCR7^low^ CD8^+^ T_EMRA_ cells (**D**) detected in COVID-19 ASYMP, SYMP and Healthy individuals. The results are representative of two independent experiments on each individual. The indicated *P* values, calculated using an unpaired t-test, show statistical significance between SYMP and Healthy individuals.

### 6. The activation status, senescence, and exhaustion profile were significantly increased in COVID-19 SYMP individuals compared to Healthy individuals within CD8^+^ T cells

Most viral infections induce activation of CD8^+^ T cells that can be detected by increases in the co-expression of CD38 and Human leukocyte antigen-DR isotype (HLA-DR). HLA-DR is constitutively expressed by antigen-presenting cells (APCs) and is involved in the presentation of antigens to T-cells. Most T-cells do not express it, but notably, a subset of activated T-cells becomes HLA-DR^+^ during an immune response. In contrast, CD38 is constitutively expressed by naive T-cells, down-regulated in resting memory cells, and then elevated again in activated cells. Thus, we evaluated the degree of CD8^+^ T-cell activation in COVID-19 positive ASYMP and SYMP patients and Healthy Controls. Within the CD8^+^ population, we analyzed markers commonly related to T cell activation (HLA-DR and CD38). Our gating strategy was used to analyze markers related to activation status, senescence, and exhaustion together within SARS-CoV-2 specific CD8^+^ T cells (**Fig. 6A)**. The levels of T-cell activation were significantly higher (hyperactivated) in SARS-CoV-2 infected SYMP patients than in healthy controls, as reflected by higher proportions of HLA-DR^+^CD38^+^ CD8^+^ T cells (**Fig. 6B**, *P* = 0.02).

**Figure 6:**
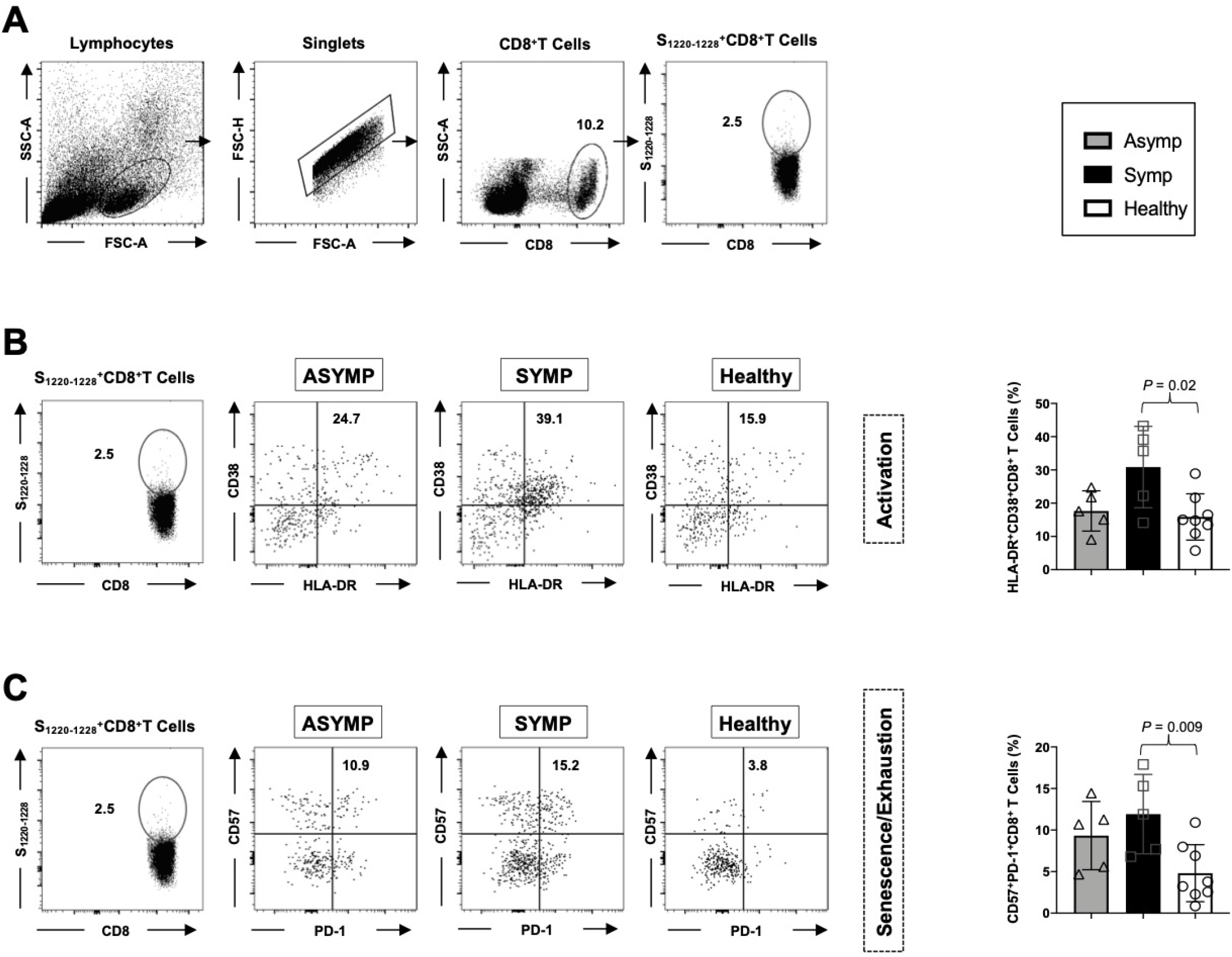
The activation status, senescence, and exhaustion profile were significantly increased in COVID-19 SYMP individuals compared to Healthy individuals within CD8^+^ T cells.Expression of CD38 and HLA-DR was detected to analyze the activation status of CD8^+^ T cells. Expression of CD57 and PD-1 was detected to analyze the senescence/exhaustion status of CD8^+^ T cells. (**A**) Gating strategy used to analyze markers related to activation status, senescence, and exhaustion together within SARS-CoV-2 specific CD8^+^ T cells. Activated cells are CD38^+^HLA-DR^+^ ; exhausted/senescent are PD1^+^CD57^+^. FACS was used to determine the expression level of various markers on tetramer gated CD8^+^ T cells specific to the S_1220-1228_ epitope. (**B**) Representative FACS data of the frequencies of HLA-DR^+^CD38^+^ CD8^+^ T cells, specific to S_1220-1228_ epitope detected in PBMCs from HLA-A*02:01 positive COVID-19 ASYMP individuals, SYMP individuals, and Healthy controls (*left panel*). Average frequencies of PBMCs-derived HLA-DR^+^CD38^+^ CD8^+^ T cells, specific to S_1220-1228_ epitope, were detected from COVID-19 ASYMP, SYMP and Healthy individuals (*right panel*). (**C**) Representative FACS data of the frequencies of CD57^+^PD-1^+^ CD8^+^ T cells, specific to S_1220-1228_ epitope, detected in PBMCs from HLA-A*02:01 positive COVID-19 ASYMP individuals, SYMP individuals, and Healthy individuals (*left panel*). Average frequencies of PBMCs-derived CD57^**+**^ PD-1^+^ CD8^**+**^ T cells, specific to the S_1220-1228_ epitope, were detected from ASYMP, SYMP and Healthy individuals (*right panel*). The results are representative of two independent experiments on each individual. The indicated *P* values, calculated using an unpaired t-test, show statistical significance between SYMP and Healthy individuals.

We evaluated the senescence/exhaustion molecules expression on circulating T cells by analyzing markers CD57 and PD-1 (**Fig. 6C**). We found that the proportion of CD57^+^PD-1^+^ CD8^+^ T cells was significantly higher in COVID-19 positive SYMP patients than in Healthy controls and ASYMP patients. Still, there was no statistical difference in the proportion of CD57^+^PD-1^+^ CD8^+^ T cells in ASYMP and SYMP patients (**Fig. 6C**, *P* = 0.009).

Taken together, findings from COVID-19 positive SYMP individuals indicate the involvement of hyperactivated CD8^+^ T cells and T cell exhaustion/senescence in the immunopathogenesis of SARS-CoV2 infection.

### 7. Elevated Plasma levels of selective cytokines in COVID-19 ASYMP and SYMP individuals compared to Healthy controls

Many studies have previously reported that hyperinflammatory response induced by SARS-CoV-2 is a major cause of disease severity and death. Therefore, we implemented a multiplex cytokine assay (Luminex) to measure inflammatory cytokines known to contribute to pathogenic inflammation (IL)-6, IL-8, tumor necrosis factor (TNF)-α, interferon (IFN)-γ and IL-17. in the plasma samples of COVID-19 in ASYMP, SYMP and Healthy individuals. The cytokines assessed in this study had different detection ranges, with IL-6 and IL-8 having the most dynamic profile followed by TNF-α, IL-17, and IFN-γ. We found that TNF-α and IFN-γ (*P* = 0.001) were significantly elevated in COVID-19 symptomatic patients compared to Healthy controls (**Fig. 7A** and **7B**). Similarly, we found that IL-6 and IL-8 (*P* = 0.01) were significantly elevated in COVID-19 symptomatic patients compared to Healthy controls (**Fig. 7C** and **7D**). Interleukin (IL)-17 is one of the many cytokines released during SARS-CoV-2 infection. IL-17 plays a crucial role in neutrophil recruitment and activation. Neutrophils subsequently can migrate to the lung and are heavily involved in the pathogenesis of COVID-19. We found that SARS-CoV-2 positive ASYMP and SYMP individuals had significantly higher levels of IL-17 than Healthy controls (*P* = 0.04) (**Fig. 7E**).

**Figure 7:**
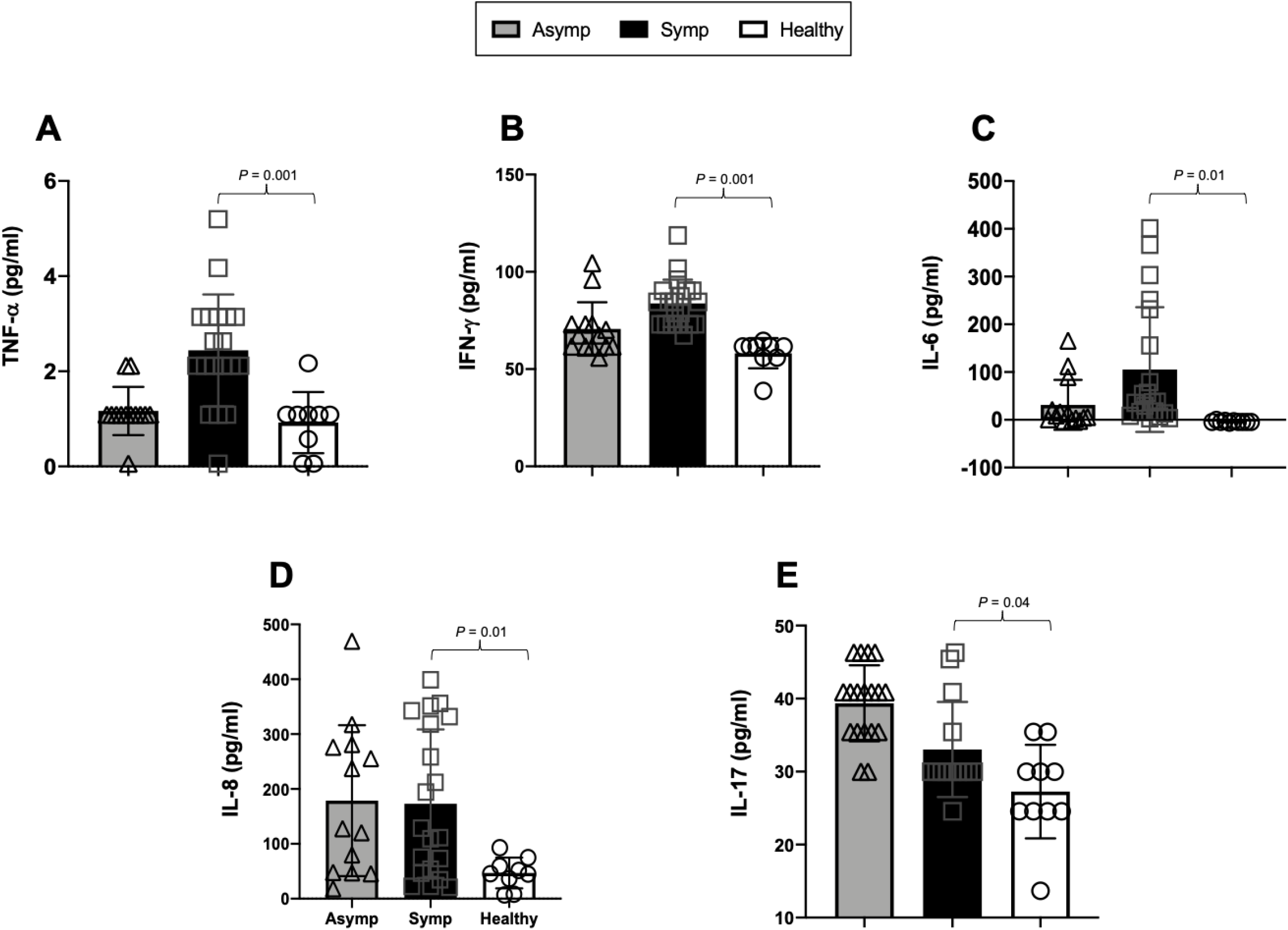
Elevated Plasma levels of selective cytokines in COVID-19 ASYMP and SYMP individuals compared to Healthy controls. Cytokine expression levels of two replicates per sample were measured in plasma samples of COVID-19 ASYMP, SYMP and Healthy individuals using Luminex. (**A**) Bar graphs with individual values showing the average amount of TNF-α (pg/ml) produced from ASYMP, SYMP and Healthy individuals. (**B**) Bar graphs with individual values showing the average amount of IFN-γ (pg/ml) produced from ASYMP, SYMP and Healthy individuals. (**C**) Bar graphs with individual values show the average IL-6 (pg/ml) produced from ASYMP, SYMP and Healthy individuals. (**D**) Bar graphs with individual values show the average IL-8 (pg/ml) produced from ASYMP, SYMP and Healthy individuals. (**E**) Bar graphs with individual values show the average IL-17 (pg/ml) produced from ASYMP, SYMP and Healthy individuals.

The vast majority of SYMP patients demonstrated elevated cytokines or cytokine storm compared to Healthy controls. In contrast, the cytokine levels were not significantly different in COVID-19 ASYMP and SYMP individuals.

Overall, our findings report that NK, CD4^+^ T cells, and CD8^+^ T cells’ phenotypic and functional characteristics in severe COVID-19 infection were compatible with activation of dysfunction/exhaustion pathways.

## DISCUSSION

COVID-19 is characterized by enhanced lymphopenia in the peripheral blood and altered T cells phenotypes shown by a spectrum of activation and exhaustion. However, antigen-specific T cell responses are emerging as a critical mechanism for both virus clearance and the most plausible pathway to long-term immunological memory that would protect against re-infection. As a result, T cell responses are of great importance in the development of vaccines (22). Moreover, post-infection changes in the composition and function of T cell subsets have significant ramifications on the patients’ long-term immunological functions (23). The impairment of effector T cell responses has been associated with the overexpression of inhibitory and senescent markers on T cells. Therefore, the main objective of this research study was to detect T-cell immune signatures in peripheral blood, including those of innate cells, and to determine how important indicators of activation and exhaustion are related to the development of symptomatic COVID-19. Factors influencing the formation and nature of protective immunity and severity of COVID-19 are still unknown. Nevertheless, data defining disease phenotypes have the prospect of informed development of new therapeutic approaches for treating individuals infected with SARS-CoV-2 and developing novel vaccines. CD57 is a marker on some cell subsets, including T cells (15, 24, 25). A costimulatory molecule like CD28 (that provides signaling for T cell activation) is expressed by naïve T cells after antigen recognition that may bind to B7 proteins to provide co-stimulatory signals (26-28). However, repeated T-cell stimulation and activation leads to gradual loss of CD28, a distinct characteristic of memory or terminally differentiated cells, and subsequent upregulation of CD57 (29-31). These senescent cells are characterized by loss of CD27 and display low proliferative capacity of the cells (32), eventually leading to an inability to eradicate infection.

The CD57 antigen is commonly used to identify populations of late-differentiated ‘senescent’ cells with defined cell phenotypes and effector functions (33, 34). In this report, we examined the patterns of expression of CD57 on NK cells and SARS-CoV-2-specific T-cells and determined increased expression of these exhaustive and senescent markers in symptomatic individuals compared to those with asymptomatic infections and Healthy controls. While CD57 is now well-recognized as a marker for terminally differentiated T-cells, it was originally thought to identify cells with natural killer activity. The expression of CD57 varies among NK cell subsets. NK cells are innate effector lymphocytes that respond to acute viral infections but might also contribute to immunopathology. NK cells are typically divided into CD56^bright^ NK cells and CD56^dim^ NK cells, which rapidly respond during diverse acute viral infections in humans, including against dengue virus, hantavirus, tick-borne encephalitis virus, and yellow fever virus, among others (35-37). Although a similar analysis of NK cells has not been performed in acute SARS-CoV-2 infection-causing COVID-19, early reports from the pandemic (in line with our findings) have indicated low circulating NK cell numbers in patients with moderate and severe disease (38-40). The SARS-CoV-2 infection has also been linked to reduced NK cell counts during the acute phase of infection. We determined a terminally differentiated phenotype with up-regulated levels of CD57 molecules in NK cells from SYMP COVID-19 patients.

CD57 is expressed by CD16^pos^ CD56^dim^ cytotoxic NK cells and CD16^pos^ CD56^neg^ inflammatory NK cells, whereas CD16^neg^ CD56^bright^ regulatory NK cells do not express this marker even during chronic infections (41-43). The acquisition of CD57 thus follows the natural differentiation of NK cells (from regulatory to cytotoxic to inflammatory NK cells). Thus, like T cells, NK cell expression of CD57 could be considered a marker of terminal differentiation, albeit not associated with senescence in this population.

The majority of prior studies into the biology of CD57 focused on the antigen’s significance in distinct T cell subsets. In the late phases of differentiation, CD57 has been found both on CD4 and CD8 T cells (15). CD57 identifies terminally differentiated cells with decreased proliferative responses in CD8 T lymphocytes. T cell senescent markers were more associated with CD8^+^ T cells than CD4^+^ T cells, consistent with our results that accumulate at lower frequencies for CD4^+^ T cells in the human periphery (44). Our findings herein indicate an increased expression of CD57^+^ T cell subsets in symptomatic patients. It was previously shown that PD-1^+^CD57^+^ CD8^+^ T cells had increased sensitivity to apoptosis mediated by PD-1 (45).

The increased expression of CD57 and PD-1 double-positive markers on CD8^+^ T cells in COVID-19 suggests that these cells are at a higher risk of apoptosis. The fraction of T cells that express CD57 increased in the symptomatic individuals, suggesting that the observed phenotypic changes may lower the T cell repertoire’s responsiveness to SARS-CoV-2 antigens, resulting in an impaired ability to eliminate the infection. CD57^+^ memory T cells accumulate in peripheral blood throughout life, especially after infection with CMV (46). These associations with age and persistent antigenic drive were mechanistically linked in an *in vitro* study, which reported that replicative senescent memory CD8^+^ T cells expressed CD57 (15). However, an earlier study had reached a different conclusion (47), and later experiments showed that CD57^+^ memory CD8^+^ T cells could proliferate *in vitro* in the presence of certain growth factors, potentially mimicking the *in vivo* microenvironment (48). Recent studies suggest that TEMRA cells are fairly resistant to apoptosis and remain in the *CD8*^*+*^ lineage for an estimated half-life of about 25 years, assuming simple exponential decline without phenotypic change (49, 50). CD8^+^ TEMRA cells that expressed CD57 were recently reported to be more sensitive to cell death than CD8^+^ TEMRA cells that lacked CD57 in response to severe stimulation with supraphysiological doses of phytohemagglutinin and interleukin-2 (51).

Compared to asymptomatic individuals and Healthy controls, symptomatic patients had increased CD57 expression on CD8^+^ TEMRA^+^ memory cells. Compared to asymptomatic infections, the phenotypic abnormality of T cells during COVID-19 infections is more apparent in symptomatic individuals and is associated with higher expression levels of exhaustive and senescent markers.

The findings of this research contribute to the current knowledge of the innate and adaptive immune landscape in asymptomatic and symptomatic COVID-19 patients. However, we recognize limitations that could be addressed with bigger sample numbers and matched control groups. Furthermore, the phenotype and activity of immune cells from the lungs performing a direct role in establishing symptomatic infections, are unknown. As a result, the immunophenotypic traits in the lungs may not completely mirror the hierarchy of immunodominant circulating immune cells in the blood.

In conclusion, this study presents an in-depth analysis of NK and T cell phenotypic and functional characteristics that are associated with COVID-19 severe disease. The finding will inform future immunotherapies to alleviate the symptoms of severe COVID-19 severe disease.

## ACKNOWLEDGEMENTS

This work is supported by the Fast-Grant PR12501 from Emergent Ventures, a grant from Herbert Family Trust, by Public Health Service Research grants AI158060, AI150091, and AI143348, AI147499, AI143326, AI138764, AI124911 and AI110902 from the National Institutes of Allergy and Infectious Diseases (NIAID) to LBM.

The authors would like to thank Dr. Dale Long from the NIH Tetramer Facility (Emory University, Atlanta, GA) for providing the Tetramers used in this study. We thank UC Irvine Center for Clinical Research (CCR) and Institute for Clinical & Translational Science (ICTS) for providing human blood samples used in this study. A special thanks to Dr. Delia F. Tifrea for her continuous efforts and dedication in providing COVID-19 samples that are crucial for this clinical research. We also thank those who contributed directly or indirectly to this COVID-19 vaccine project: Dr. Steven A. Goldstein, Dr. Michael J. Stamos, Dr. Suzanne B. Sandmeyer, Jim Mazzo, Dr. Daniela Bota, Dr. Beverly L. Alger, Dr. Dan Forthal, Dr. Tahseen Muzaffar, Dr. Ilhem Messaoudi, Anju Subba, Janice Briggs, Marge Brannon, Beverley Alberola, Jessica Sheldon, Rosie Magallon, and Andria Pontello.

